# Context-dependent miR-21 regulation of TLR7-mediated autoimmune and foreign antigen driven antibody-forming cell and germinal center responses

**DOI:** 10.1101/2021.03.12.435182

**Authors:** Stephanie L. Schell, Kristen N. Bricker, Adam J. Fike, Sathi Babu Chodisetti, Phillip P. Domeier, Nicholas M. Choi, Melinda J. Fasnacht, Sara A. Luckenbill, Steven F. Ziegler, Ziaur S.M. Rahman

**Affiliations:** Department of Microbiology and Immunology, Pennsylvania State University College of Medicine, Hershey, PA 17033; Immunology Program, Benaroya Research Institute, Seattle, WA 98101

## Abstract

MicroRNAs (miRNAs) are involved in healthy B cell responses and the loss of tolerance in systemic lupus erythematosus (SLE), though the role of many miRNAs remains poorly understood. Dampening miR-21 activity was previously shown to reduce splenomegaly and blood urea nitrogen levels in SLE-prone mice, but the detailed cellular responses and mechanism of action remains unexplored. Herein, using the TLR7 agonist imiquimod-induced SLE model, we observed that loss of miR-21 in *Sle1b* mice prevented the formation of plasma cells and autoantibody forming cells (AFC), without a significant effect on the magnitude of the germinal center (GC) response. We further observed reduced dendritic cell and monocyte numbers in the spleens of miR-21 deficient *Sle1b* (*Sle1b*.miR-21KO) mice that were associated with reduced interferon, proinflammatory cytokines, and effector CD4^+^ T cell responses. RNAseq analysis on B cells from *Sle1b*.miR-21KO mice revealed reduced activation and response to interferon and cytokine and target array analysis revealed modulation of numerous miR-21 target genes in response to TLR7 activation and type I interferon stimulation. Our findings in the B6.*Sle1b*.Yaa spontaneous model recapitulated the miR-21 role in TLR7-induced responses with an additional role in autoimmune GC and Tfh responses. Finally, immunization with T-dependent antigen revealed a role for miR-21 in foreign antigen driven GC and Ab, but not AFC responses. Our data suggest a potential multifaceted, context-dependent role for miR-21 in autoimmune and foreign antigen driven AFC and GC responses. Further study is warranted to delineate the cell-intrinsic requirements and mechanisms of miR-21 during infection and SLE development.

**Key Points:**

- miR-21 has context dependent effects on AFC and GC responses
- miR-21 promotes TLR7-driven autoimmunity with activation of multiple B cell pathways
- miR-21 is required for optimal B cell responses to T-dependent foreign antigen

## Introduction

Systemic lupus erythematosus (SLE) is one of over 100 currently identified autoimmune diseases. SLE patients develop multisystemic organ manifestations due to immune complex deposition and recruitment of myeloid cells to local tissues, and systemic inflammation following the loss of B cell tolerance (1). Morbidity and mortality in SLE patients is primarily associated with the development of lupus nephritis or cardiovascular events (2, 3).

Loss of B cell tolerance in SLE is well-documented in the secondary lymphoid organs, with aberrant selection occurring at multiple stages in the spleen (4–9). This aberrant selection of B cells generates long-lived, class-switched autoantibody-secreting B cells in SLE patients that are the target of current therapeutic strategies for SLE. Studies focused on understanding the loss of tolerance in SLE have highlighted the role of TLR signaling, interferon, and cytokine responses in driving SLE initiation and progression (10–16). Despite these advances, there is currently only one FDA approved therapy for SLE, belimumab, which broadly targets B cell survival via blocking B cell access to BAFF (17). Belimumab exhibits moderate efficacy for SLE, but at the cost of reducing anti-pathogen immunity, leaving room for substantial improvement of therapeutic strategies that may preserve anti-pathogen responses while improving quality of life in SLE patients. The development of novel therapeutics for SLE relies on better understanding of the mechanisms and additional factors underlying disease development and progression.

microRNAs (miRNAs) have been implicated in the control of B cell functions, including development and response to foreign antigen (18–24). As such, it is not surprising that the miR-17∼92 cluster, miR-155, miR-146a, and miR-148a also have well-delineated roles in the establishment or restraint of SLE and related autoimmune diseases (25–30). It is likely that additional miRNAs are involved in SLE development and/or progression. Studying miRNA function in SLE is attractive since miRNAs have been a focus of recent therapeutic approaches for other diseases, including hepatitis C viral infection and cancer (31). While miRNA focused therapeutics have not yet been developed specifically for the treatment of autoimmune diseases, they present an intriguing possibility for directed targeting in the future.

miR-21 expression increases in SLE-prone mice and SLE patients (32–38). Additionally, a previous study using an SLE mouse model observed reduced blood urea nitrogen and splenomegaly when miR-21 was antagonized (32). Specific mechanisms, including autoreactive B cell development, GC responses, AFC generation, cellular subset identification, and potential target genes remain unclear, prompting further investigation of the mechanisms underlying the role of miR-21 in the development of SLE. As such, we assessed the role of miR-21 in imiquimod (IMQ)-treated autoimmune-prone *Sle1b* mice deficient for miR-21, a TLR7-induced SLE model previously established for the study of robust B cell and SLE responses (39, 40). We also examined the role of miR-21 in autoimmune AFC, GC and Tfh responses in a TLR7-overexpressing B6.*Sle1b*.Yaa (*Sle1b*^Yaa^) spontaneous SLE model. In addition to the unclear role of miR-21 in SLE-associated AFC and GC responses, the impact of miR-21 on AFC and GC responses to foreign antigen remained unexplored, which we also sought to characterize.

Our results from the TLR7-induced SLE model reveal that the increased number of innate immune cells contributed to TLR7 and miR-21 driven splenomegaly in *Sle1b* mice. miR-21 promoted splenic autoreactive AFC formation, and population of autoreactive AFCs in the bone marrow in IMQ-treated *Sle1b* mice. The onset of autoimmunity was linked to reduced plasma cell responses, but not germinal center formation, indicating that miR-21 predominantly affects plasma cell formation in this model. As such, interferon, pro-inflammatory cytokines, and the B cell survival factor, BAFF were reduced in the spleens of IMQ-treated *Sle1b*.miR-21KO mice. Reduced cytokine responses were associated with reduced DC and monocyte infiltration and reduced effector CD4^+^ T cell responses. B cells in *Sle1b*.miR-21KO mice were less activated and pathway analysis revealed reduced response to pro-inflammatory cytokines and interferon and reduced activation signature. Our findings in the Sle1b^Yaa^ spontaneous model recapitulated the role of miR-21 in regulating splenomegaly, innate cell numbers, AFC/plasma cell responses and autoantibody production with the additional requirement of miR-21 in autoimmune GC and Tfh responses. These data collectively suggest that miR-21 has the capacity to act in multiple immune cell subsets in SLE, including direct involvement in B cell activation, modulating effector T cell responses, and the recruitment and activation of myeloid cells, which may drive autoimmune AFC and GC generation and systemic autoimmunity development, in a model dependent manner. Interestingly, we also observed an effect of miR-21 on responses to TD-antigen following immunization which affect GC, Tfh and antigen specific Ab responses, but not overall AFC or plasma cell response. These data together indicate an important context-dependent miR-21 role in regulating autoimmune and foreign antigen driven AFC and GC responses.

## Materials and Methods

### Mice

For autoimmunity experiments, *Sle1b* mice [detailed previously (41)] were crossed to miR-21KO mice (Jackson Laboratory stock #016856) to generate *Sle1b*.miR-21KO. Only female mice were used for autoimmunity experiments involving imiquimod treatment of mice due to the greater penetrance of autoimmunity in females. In autoimmunity experiments involving the use of TLR7-overexpressing B6.*Sle1b*.Yaa mice and B6.*Sle1b*.Yaa mice deficient in miR-21, only male mice were used as they carry the Yaa locus. For immunization experiments, wild-type B6/129F2/J (B6.129F2) (stock #101045) and miR-21KO female and male mice were used. All mice were housed in a specific pathogen free facility and all experiments were performed under the guidelines of the Pennsylvania State University Institutional Animal Care and Use Committee.

### Immunization of Mice

7-9 week old mice were immunized with 4-hydroxy-3-nitrophenol–Keyhole Limpet Hemocyanin (NP-KLH) (Biosearch Technologies). NP-KLH was premixed in a 1:1 mixture of PBS: Complete Freund’s Adjuvant (CFA) (Sigma Aldrich) prior to injection. Mice received 200μg of NP-KLH in 200μl of injection cocktail i.p. on d0 and were harvested for analysis at the indicated time points post-immunization.

### Epicutaneous Imiquimod Treatment

Imiquad 5% imiquimod cream (Glenmark Pharmaceuticals) was applied to the back surface of both mouse ears beginning at 8-10 weeks of age for 8-12 weeks, 3x per week to study autoimmunity as previously described (39, 42).

### Flow Cytometry and Antibodies

In brief, spleens were prepared for flow cytometry by generating single cell suspensions. RBCs were lysed by Tris Ammonium Chloride. Single cell suspensions of bone marrow cells were prepared by flushing the marrow followed by RBC lysis. Resulting single cell suspensions of splenocytes or bone marrow cells were Fc blocked with Trustain FcX (anti-mouse CD16/32 clone 93-Biolegend) and stained with combinations of the following antibodies: anti-B220-PacBlue (RA3-6B2), GL7-FITC, anti-CD95-PeCy7 (Jo2), anti-CD4-Alexa Flour 700 (RM4-5), anti-CXCR5-biotin (2G8), anti-PD-1-PE (29F.1A12), anti-CD11c-BV421 (N418), anti-NK1.1-biotin (PK136), anti-Ly6G-BV711 (1A8), CD86-PeCy5 (GL-1), anti-CD44-BV605/APC (IM7), anti-CD62L-PeCy7 (MEL-14), anti-CD80-PE (16-10A1), anti-CD19-BV605 (6D5), anti-CD267-PE (8F10), anti-CD138-PeCy7 (281–2), and SA-PeCy5 from Biolegend; anti-CD43-FITC (S7), anti-Ly6C-FITC (AL-21), anti-CD11b-AF700 (M1/70), anti-CD90.2-biotin (53-2.1), anti-CD19-biotin (1D3), anti-CD40-FITC (HM40-3), anti-IgD-APC (11-26c.28), and SA-V500 from BD Biosciences. Viability staining was performed by incubating samples with Fixable Viability Dye (e780) from eBioscience. Data were acquired on an LSRII flow cytometer (BD Biosciences) and data were analyzed using FlowJo software (Tree Star).

### Immunohistology and Antibodies

Spleens were flash-frozen in OCT compound. 6μm sections were cut on a cryostat and fixed in cold acetone as previously described (43). Spleen sections were stained with the following combination of antibodies: GL7-FITC, anti-CD4-PE (GK1.5), anti-IgD-APC (11-26c.2a) from BD Biosciences. Image acquisition was performed on a Leica DM4000 fluorescent microscope. The color intensity of images was consistently enhanced among images using Adobe Photoshop for better visualization, while maintaining the integrity of the data. For GC quantitation, GC size was measured on 10 randomly selected GCs (or the highest number available on the section) from 4 or more mice per group.

### ELISA

For NP-specific ELISAs, plates were coated with NP29-BSA (low affinity) or NP4-BSA (high affinity) (Biosearch Technologies). For ANA ELISAs, plates were coated with poly-L-lysine (Sigma Aldrich) and then subsequently coated with salmon sperm dsDNA (Invitrogen) or Smith ribonucleoprotein (SmRNP) (Arotec Diagnostics). Nucleosome plates were generated by coating with dsDNA followed by histone (Sigma Aldrich). For detection, biotinylated anti-IgG (Jackson Immunoresearch), anti-IgG1, anti-IgG2a, anti-IgG2b, or anti-IgG2c (Southern Biotech), followed by streptavidin-alkaline phosphatase (Vector Laboratories) were used for the detection of respective subtypes. Development was performed with PNPP (*p*-nitrophenyl phosphate, disodium salt) substrate (Thermo Fisher). Quantitation was performed as previously described (44).

### RNA isolation, cDNA synthesis, and qPCR

Cells were lysed with Trizol reagent as indicated by the manufacturer’s protocol (Ambion). Following chloroform extraction and aqueous phase recovery, an equal volume of 70% ethanol was added to the isolated aqueous phase, mixed, and loaded onto RNeasy columns (Qiagen) for further purification. On column DNase digestion (Qiagen) was performed to eliminate gDNA and the remainder of the standard RNeasy column purification protocol was performed thereafter. cDNA was synthesized from isolated RNA using the Applied Biosystems High Capacity cDNA Reverse Transcription Kit. qPCR was performed using Integrated DNA Technologies (IDT) pre-designed Primetime assay probes for the following genes: IL-10 (Mm.PT.58.13531087), IL-6 (Mm.PT.58.10005566), TNF (Mm.PT.58.12575861), BAFF (Mm.PT.58.29965375), IL-21 (Mm.PT.58.43546164), IFNγ F (5’ CGG CAC AGT CAT TGA AAG CC 3’) and R (5’ TGC ATC CTT TTT CGC CTT GC 3’), IL-12p35 F (5’ CAC AAG AAC GAG AGT TGC CTG GCT 3’ and R (5’ GGT CTG CTT CTC CCA CAG GAG GTT 3’), and HPRT (Mm.PT.39a.22214828). Gene expression was normalized to HPRT. qPCR was performed using Power Sybr Green Master Mix (Thermo Fisher) and cycling was performed as follows: 10 mins at 95°C, then 40 cycles at 95°C for 15s and 62°C for 1min on a Step One Plus Instrument (Applied Biosystems). All qPCR was performed in technical triplicate and the average of these replicates was used to generate one data point per mouse assessed, as indicated in the figure legends. Fold change was calculated by the 2^-ΔΔCT^ method.

### miRNA Isolation, cDNA synthesis, and qPCR

Isolated cells pooled from multiple mice per group were processed using an Invitrogen mirVana miRNA Isolation kit (Invitrogen) according to manufacturer’s instructions. The small RNA fraction, including miRNAs, was used for cDNA synthesis according to the Taqman MicroRNA Reverse Transcription Kit protocol (Applied Biosystems). Taqman miRNA assays for the housekeeping control gene U6 snRNA (#001973) and hsa-miR-21-5p (which also recognizes mmu-miR-21a-5p) (#000397) were purchased from Thermo Fisher. qPCR was performed using Taqman Universal Master Mix II, with UNG (Thermo Fisher). Samples were run on a Step One Plus instrument (Appled Biosystems) according to the following cycling protocol: 2 mins at 50°C, followed by 10 mins at 95°C, followed by 40 cycles of 95°C for 15s then 60°C for 1 min. All qPCR was performed in technical triplicate. Fold change was calculated by the 2^-ΔΔCT^ method.

### RNA Seq Sample Preparation and Analysis

Total B cells were isolated from spleens of *Sle1b* or *Sle1b*.miR21KO mice after 8 weeks of topical imiquimod treatment by negative selection using the mouse B cell isolation kit from STEMCELLTechnologies. Total RNA was prepared from 5 x10^6^ purified cells per mouse using the protocol detailed in the *RNA isolation, cDNA synthesis, and qPCR* section. Total RNA (0.5ng/mL) was added to lysis buffer from the SMART-seq V4 Ultra Low Input RNA kit for Sequencing (Takara), and reverse transcription was performed followed by PCR amplification to generate full-length amplified cDNA. Sequencing libraries were constructed using the NexteraXT DNA sample preparation kit (Illumina) to generate Illumina-compatible barcoded libraries. Libraries were pooled and quantified using a Qubit^®^ Fluorometer (Life Technologies). Dual-index single-read sequencing of pooled libraries was carried out on a HiSeq2500 sequencer (Illumina) with 58-base reads using HiSeq v4 Cluster and SBS kits (Illumina) with a target depth of 5 million reads per sample. Base calls were processed to FASTQs on BaseSpace (Illumina), and a base call quality-trimming step was applied to remove low-confidence base calls from the ends of reads. The reference genome, using STAR v.2.3.2a and gene counts were generated using htseq-count. QC and metrics analysis was performed using the Picard family of tools (V1.134). Gene set enrichment plots were generated from total gene counts using GSEA (version 4.0.3), and the Principle Component Analysis was performed using R (Version 1.1.456). Differentially-expressed genes between *Sle1b* or *Sle1b*.miR21KO samples were calculated with Limma (Version 3.42.2) using a fold-change cutoff of 1.5 and an adjusted p-value cutoff of 0.05. Volcano plots were generated using the ggplot2 package in R. Pathway Analysis was performed from lists of differentially-expressed genes using GO Pathway analysis tool (45, 46) and heatmap plots were generated from differentially expressed gene lists using Java Treeview (Version 1.1.6r4).

### miR-21 Target qPCR Array

For initial characterization of miR-21 expression after stimulation, B cells were purified from the spleens of *Sle1b* mice using the Stem Cell Technologies mouse B cell isolation kit. B cells were stimulated with different combinations of R848 (100ng/mL), IFNβ (500 units/mL), and IFNα4 (500 units/mL) for the indicated time points. For the qPCR array, B cells isolated from *Sle1b* and *Sle1b*.miR-21KO mice were stimulated with R848 and IFNβ at the above concentrations for 48 hours. RNA was isolated from purified B cells as described in *RNA Isolation*. cDNA was synthesized with the first strand kit from Qiagen, with a starting input of 800ng of RNA per sample, according to manufacturer’s instructions. Qiagen mouse miR-21 target array plates (cat# PAMM-6001ZC) were used according to manufacturer’s instructions, with n=3/condition. Ct values were uploaded to Qiagen’s data analysis center and results were normalized to housekeeping gene expression among samples.

### ANA ELIspots

Multiscreen Filter plates (Millipore) were coated with Poly-L-lysine (Sigma Aldrich), and subsequently with salmon sperm dsDNA (Invitrogen) or SmRNP (Arotec Diagnostics). Nucleosome plates were generated via incubation with dsDNA followed by histone (Sigma Aldrich). Plates were blocked in 5% FCS in PBS and then plated with 1 million splenocytes or bone marrow cells for serial dilution. Detection was performed by incubation with anti-IgG-biotin (Jackson Immunoresearch) followed by streptavidin-alkaline phosphatase (Vector Laboratories). Development was performed with Vector Blue substrate (Vector Labs).

### HEp-2 ANA Assay

HEp-2 slides (Antibodies Incorporated) were used according to manufacturer’s protocol with serum diluted 1:50 in 2% BSA in PBS. Binding was visualized via incubation with anti-kappa-FITC (H139-52.1) (Southern Biotech). Imaging was performed with a Leica DM4000 fluorescent scope.

### Statistics

Multiple group comparison was performed by ANOVA with multiple comparisons analysis, Tukey test. Two group comparison was performed by Mann-Whitney analysis. Statistical significance is assigned according to following values: *p < 0.05, **p < 0.01, ***p < 0.001, and ****p < 0.0001. All statistical analysis was performed in Graph Pad Prism 6 software.

### Data Availability

The RNAseq data generated for this study is deposited in GSE148285.

## RESULTS

### IMQ-Treated Sle1b Mice Exhibit High Levels of miR-21 Expression and Loss of miR-21 Reduces Splenomegaly and Innate Cell Numbers

Previously, miR-21 inhibition was shown to reduce splenomegaly in SLE-prone mice that spontaneously develop disease (32). However, the mechanism by which miR-21 causes increased splenic cellularity, the cell types that contribute to splenomegaly and how tolerance is lost remained unclear. To test the requirement for miR-21 in these responses, we used a TLR7-induced model in which *Sle1b* mice were treated epicutaneously with the TLR7 ligand, imiquimod (IMQ, Fig. 1A). IMQ treatment enhances AFC and GC responses and accelerates SLE manifestations (40), which is in line with previous studies establishing TLR7 activation as an important driver of the loss of tolerance in SLE (10, 11, 14). Of note, similar to humans, disease development in the *Sle1b* strain is female biased (47), and therefore all studies in the IMQ model using this strain were performed solely on female mice. Importantly, treatment with IMQ on the ear surface does not cause any visible damage or swelling of the ear, eliminating the concern of pathogen introduction through wounded tissue (Fig. 1B).

**Figure 1.**
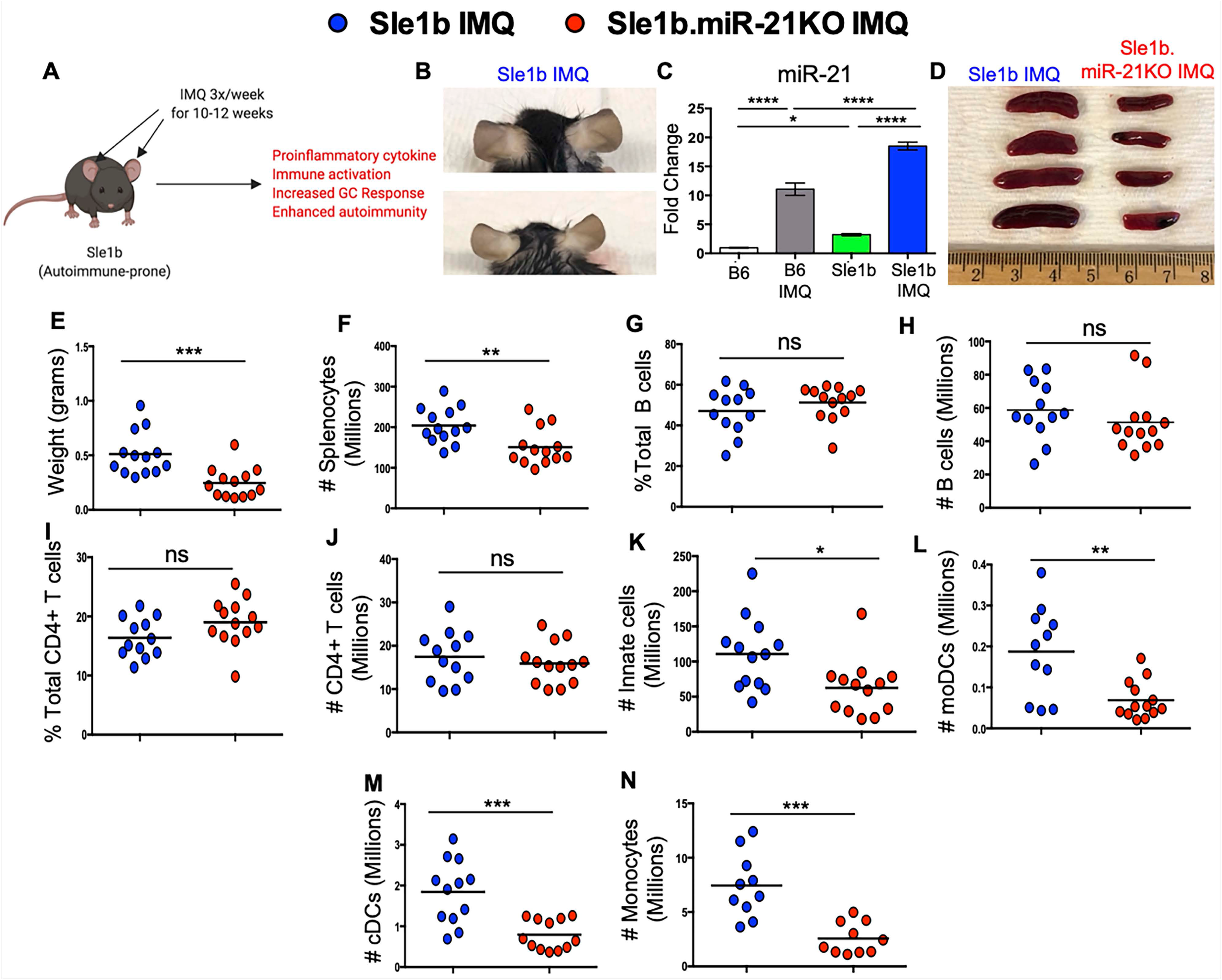
IMQ Treatment Drives miR-21 Expression and miR-21-Dependent Splenomegaly and Innate Cell Numbers in Autoimmune-Prone Sle1b Mice. All data are from *Sle1b* and *Sle1b*.miR-21KO mice following 8-12 weeks of IMQ treatment. Data are from 2 independent experiments, each with 5-8 mice per group. Each data point represents one mouse. Blue circles are *Sle1b* IMQ and red circles are *Sle1b*.miR-21KO IMQ. **(A)** Schematic illustrating the IMQ treatment model whereby mice are treated epicutaneously on the ear surface with a cream containing 5% IMQ. **(B)** Images depicting a lack of visible inflammation on the ear surface of autoimmune-prone mice treated with IMQ. **(C)** qPCR for miR-21 expression in the spleens of B6 (clear), B6 IMQ (gray), *Sle1b* (green), and Sle1b IMQ (blue) mice. Bars represent the SD of data. **(D)** Images of spleen size in the indicated strains. **(E)** Weight of spleens derived from the indicated mice. **(F)** Quantification of the number of total splenocytes. **(G-H)** Quantification of the **(G)** percentage and **(H)** number of B cells within total splenocytes. **(I-J)** Quantification of the **(I)** percentage and **(J)** number of CD4 T cells within total splenocytes. **(K-N)** Quantification of the total number of **(K)** innate cells, **(L)** moDCs, **(M)** cDCs and **(N)** monocytes in the spleen. Except for **(C)**, Bars on graphs represent the mean of the data points. Except for **(C)**, two group comparison was performed by Mann-Whitney analysis. For **(C)**, multiple group comparison was performed by one-way ANOVA with multiple comparisons analysis, Tukey test. *p < 0.05, **p < 0.01, ***p < 0.001, and ****p < 0.0001.

We first asked if miR-21 was increased in our autoimmune-prone strain, *Sle1b*, relative to wild-type controls, and if TLR7 activation could further upregulate miR-21 expression. We found that *Sle1b* mice had spontaneous increases in miR-21 expression in splenocytes, but miR-21 expression was even greater following IMQ treatment (Fig. 1C). To assess if miR-21 affects responses in *Sle1b* mice, *Sle1b* and *Sle1b*.miR-21KO mice were treated with IMQ for 12 weeks. Consistent with our published data (40), we observed large spleens in IMQ-treated *Sle1b* mice, which were significantly reduced in size and weight in IMQ-treated *Sle1b*.miR-21KO mice (Fig. 1D-E). Accordingly, the total number of splenocytes was significantly reduced in the absence of miR-21 (Fig. 1F). As the spleens of IMQ-treated *Sle1b*.miR-21KO mice were reduced in size, we asked if the number of total B cells and CD4^+^ T cells in the spleen was altered. We did not find a difference in the magnitude of these cell types in the spleen (Fig. 1G-J). We, however, found a difference in the magnitude of the innate cell response as we observed reduced numbers of innate cells including monocytes, monocyte derived (moDC), and conventional (cDC) DCs (Fig. 1K-N). These results suggested that the SLE-associated splenomegaly, likely caused by increased recruitment of innate cells driven by TLR7 activation, is dependent on miR-21 expression.

### miR-21 Controls Plasma Cell Formation, but Not GC Responses, in IMQ-treated Sle1b Mice

While the total number of B cells in the spleens of *Sle1b*.miR-21KO mice remained unchanged, we investigated whether the generation of plasma cells was altered in *Sle1b*.miR-21KO mice. We found that *Sle1b*.miR-21KO mice exhibited reduced formation of early and mature plasma cells, but not plasmablasts (Fig. 2A-D). Since GC responses are also established drivers of autoimmunity in *Sle1b* mice (40, 47) we next determined whether the magnitude of the GC and Tfh responses differed between IMQ treated *Sle1b* and *Sle1b*.miR-21KO mice. Surprisingly, we found no significant reduction in the magnitude of the GC or Tfh responses in *Sle1b*.miR-21KO mice (Fig. 2E-K). These data indicate that miR-21 does not overtly affect the GC response in IMQ treated *Sle1b* mice but significantly affects plasma cell responses.

**Figure 2.**
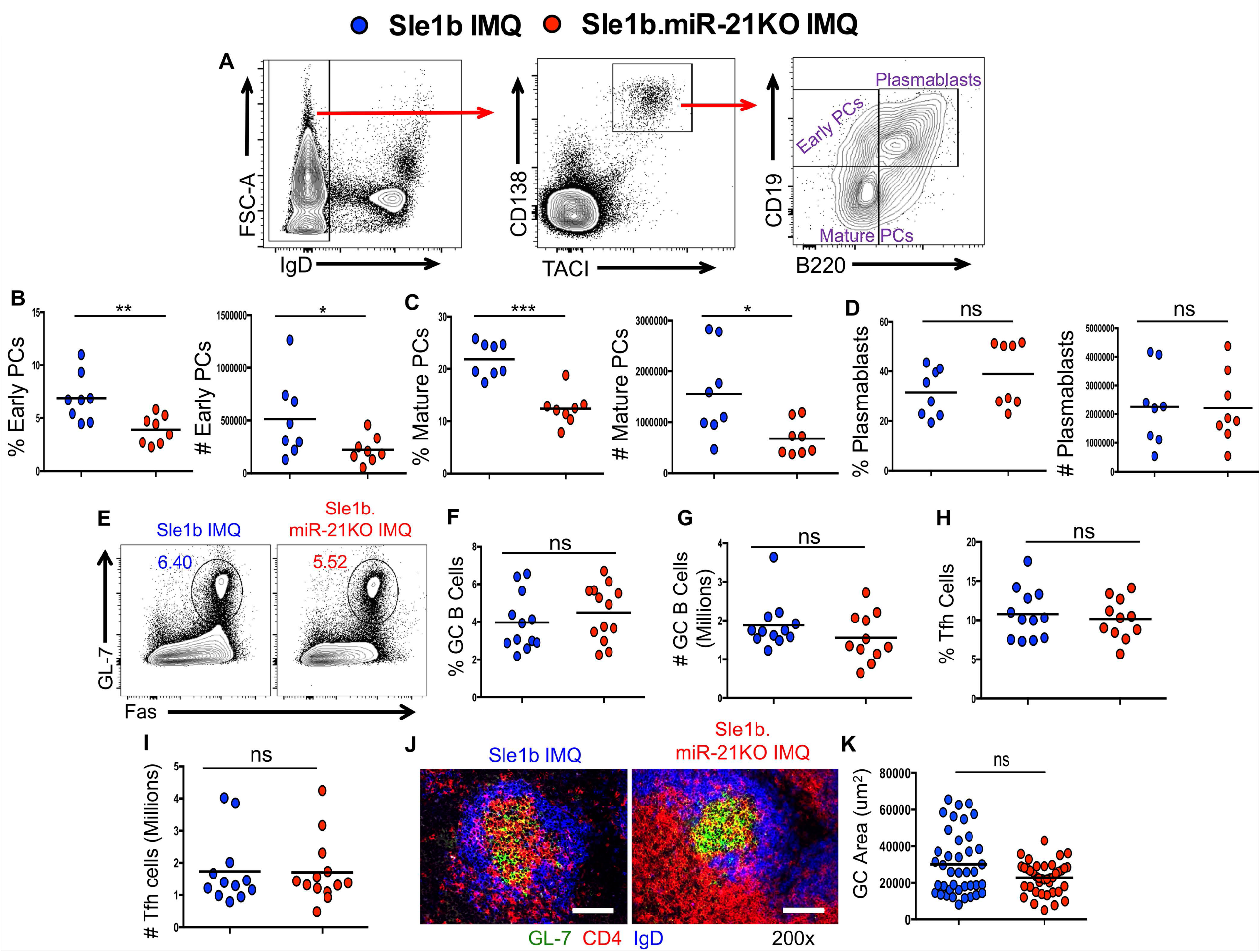
miR-21 Alters the Magnitude of the Plasma Cell but not GC Response in IMQ Treated Autoimmune-Prone Sle1b Mice. All data are from *Sle1b* and *Sle1b*.miR-21KO mice following 8-12 weeks of IMQ treatment. Data are from 2 independent experiments, each with 5-8 mice per group. Except for histology, each data point represents one mouse (biological replicate). See “Materials and Methods” for more information about histology quantitation. Blue circles are *Sle1b* IMQ and red circles are *Sle1b*.miR-21KO IMQ. **(A)** Gating strategy for identification of plasmablasts, early plasma cells, and mature plasma cells in the spleen. **(B-D)** Quantification of the percentage and numbers of **(B)** early plasma cells, **(C)** mature plasma cells, and **(D)** plasmablasts. **(E)** Representative flow plots of the GC B cell (GL-7^hi^Fas^hi^) response. **(F-G)** Quantification of the **(F)** percentage GC B cells within the total B220^+^ population and the **(G)** total number of GC B cells. **(H-I)** Quantification of the **(H)** percentage and the **(I)** total number of Tfh. **(J)** Representative images of spleen sections stained for GL-7 (green), CD4 (red), and IgD (blue). Scale bars represent 1001m. **(K)** Quantification of GC responses by measuring the GC area. Data represent at least 4 mice per genotype. Two group comparison was performed by Mann-Whitney analysis. *p < 0.05, **p < 0.01, ***p < 0.001, and ****p < 0.0001.

### miR-21 Drives the Development of Autoreactive AFCs in IMQ-Treated Sle1b Mice

After observing differences in the generation of plasma cell responses between IMQ-treated *Sle1b* and *Sle1b*.miR-21KO mice, we asked whether the fraction of B cells that secrete autoreactive antibodies against common SLE autoantigens including dsDNA and SmRNP differed between the two strains. To quantify the number of autoreactive AFCs in these mice, we performed ELISpot analysis on cells derived from the spleen and the bone marrow, the latter of which is a site of plasma cell migration following exit from the secondary lymphoid organs. Importantly, we found a drastic reduction in the number of AFCs producing IgG antibodies against dsDNA and SmRNP in the spleen (Fig. 3A) and bone marrow (Fig. 3B) in IMQ-treated *Sle1b*.miR-21KO mice compared to IMQ-treated *Sle1b* mice. This correlated with absent or low seropositivity in IMQ-treated *Sle1b*.miR-21KO mice compared to the strong seropositivity found in IMQ-treated *Sle1b* mice (Fig. 3C-D) as well as vastly decreased IgG^+^ autoantibodies of these specificities in the serum (Fig. 3E-G). Our results indicate that miR-21 affects the generation of autoreactive AFCs and as a result, autoreactive antibody titers in the serum.

**Figure 3.**
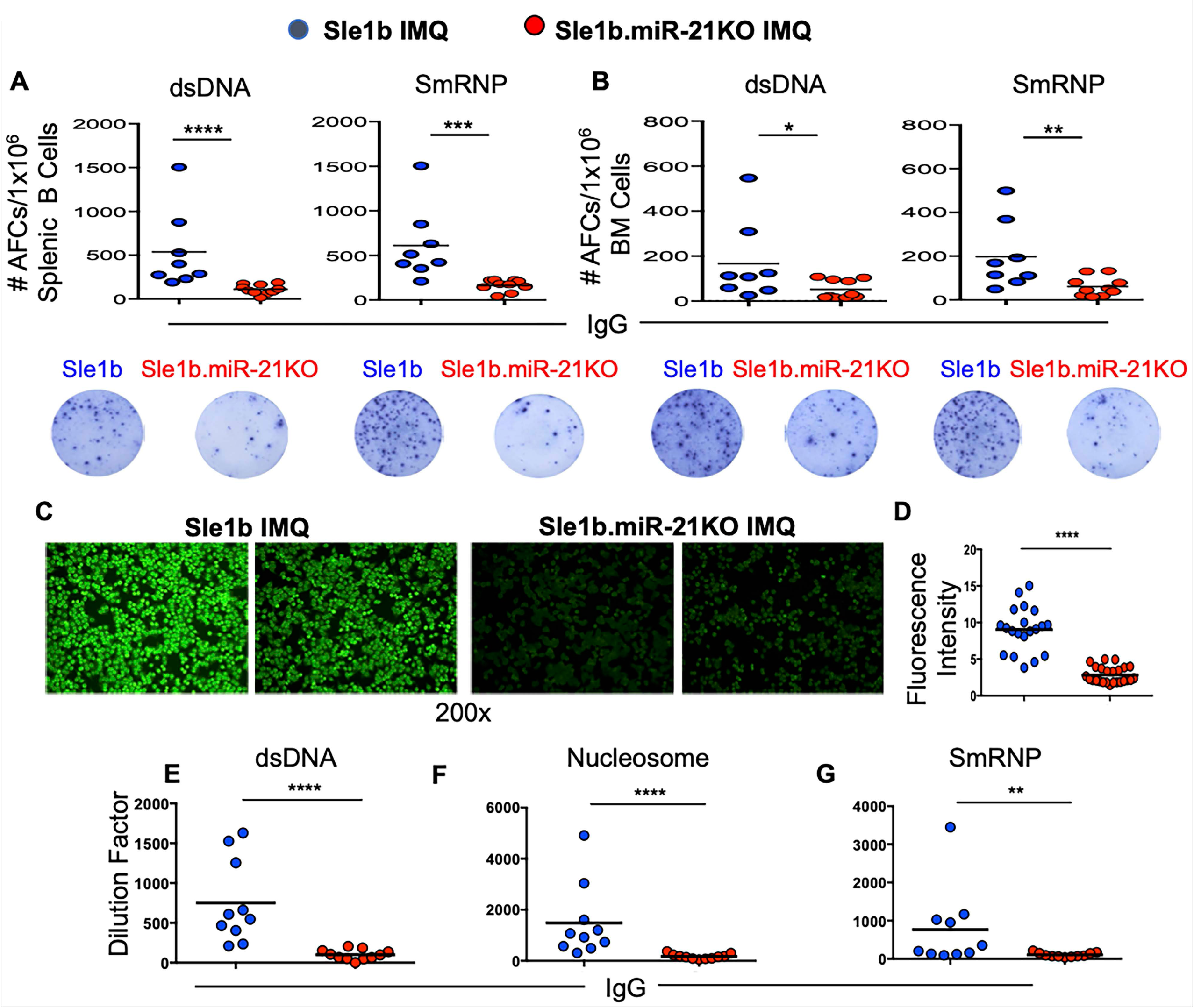
miR-21 Drives Autoreactive AFC and Autoantibody Responses. All data are from *Sle1b* and *Sle1b*.miR-21KO mice following 8-12 weeks of IMQ treatment. Blue circles are *Sle1b* IMQ and red circles are *Sle1b*.miR-21KO IMQ. Results are from 2 independent experiments, each with 4 or more mice per group. Each data point represents one mouse, except for HEp-2 quantitation which is detailed below. **(A)** Quantitation and representative images of ELISpot analysis of autoantibody secreting B cells producing antibodies with reactivity against dsDNA and SmRNP in the spleens of the indicated genotypes of mice. **(B)** Quantification and representative images of ELISpot analysis of autoantibody secreting B cells producing antibodies with reactivity against dsDNA and SmRNP in the bone marrow of the indicated genotypes of mice. **(C)** Representative images of seropositivity/seronegativity following incubation of serum on HEp-2 slides. **(D)** Quantification of the fluorescence emitted from the detection of autoreactive antibodies bound to HEp-2 cells. Staining was quantitated from 4-5 mice per group. Data points represent measurements taken from random individual 10x fields. **(E-G)** Serum titers of autoreactive antibodies reactive against **(E)** dsDNA, **(F)** nucleosome, and **(G)** SmRNP. Two group comparison was performed by Mann-Whitney analysis. *p < 0.05, **p < 0.01, ***p < 0.001, and ****p < 0.0001.

### miR-21 Drives CD4^+^ T Cell Activation and Interferon and Pro-Inflammatory Cytokine Response

Since miR-21 affects DC numbers in the spleens of IMQ-treated *Sle1b* mice, we next asked whether this is associated with CD4^+^ T cell effector responses and inflammatory cytokine and interferon levels, hallmarks of SLE development. As such, IMQ treated *Sle1b*.miR-21KO mice had a significant reduction in the percentage and number of CD4^+^CD62L^lo^CD44^hi^ effector T cells (Fig. 4A-C). Given reduced number of DCs, monocytes and CD4^+^ effector T cells in the absence of miR-21, we next determined the role of miR-21 in cytokine production in IMQ-treated *Sle1b* mice. We found significant reductions in the transcript levels of the SLE-associated cytokines *Tnfsf13b* (BAFF), *Ifng* (IFNγ), *Il6* (IL-6), and *Il10* (IL-10) in splenic tissues of *Sle1b*.miR-21KO mice compared to *Sle1b* mice (Fig. 4D). No significant reductions in *Tnf* (TNF), *Il12a* (IL-12p35), or *Il21* (IL-21) were observed (Fig. 4D). These results indicated that T cell activation and inflammatory responses are altered in *Sle1b*.miR-21KO mice. Further, the altered interferons and cytokines have well-established roles in altering autoreactive B cell generation, selection, or survival in SLE (12, 13, 15, 48). Altogether, these results support a role for miR-21 in promoting a proinflammatory cytokine environment in the spleen that can alter B cell activation.

**Figure 4.**
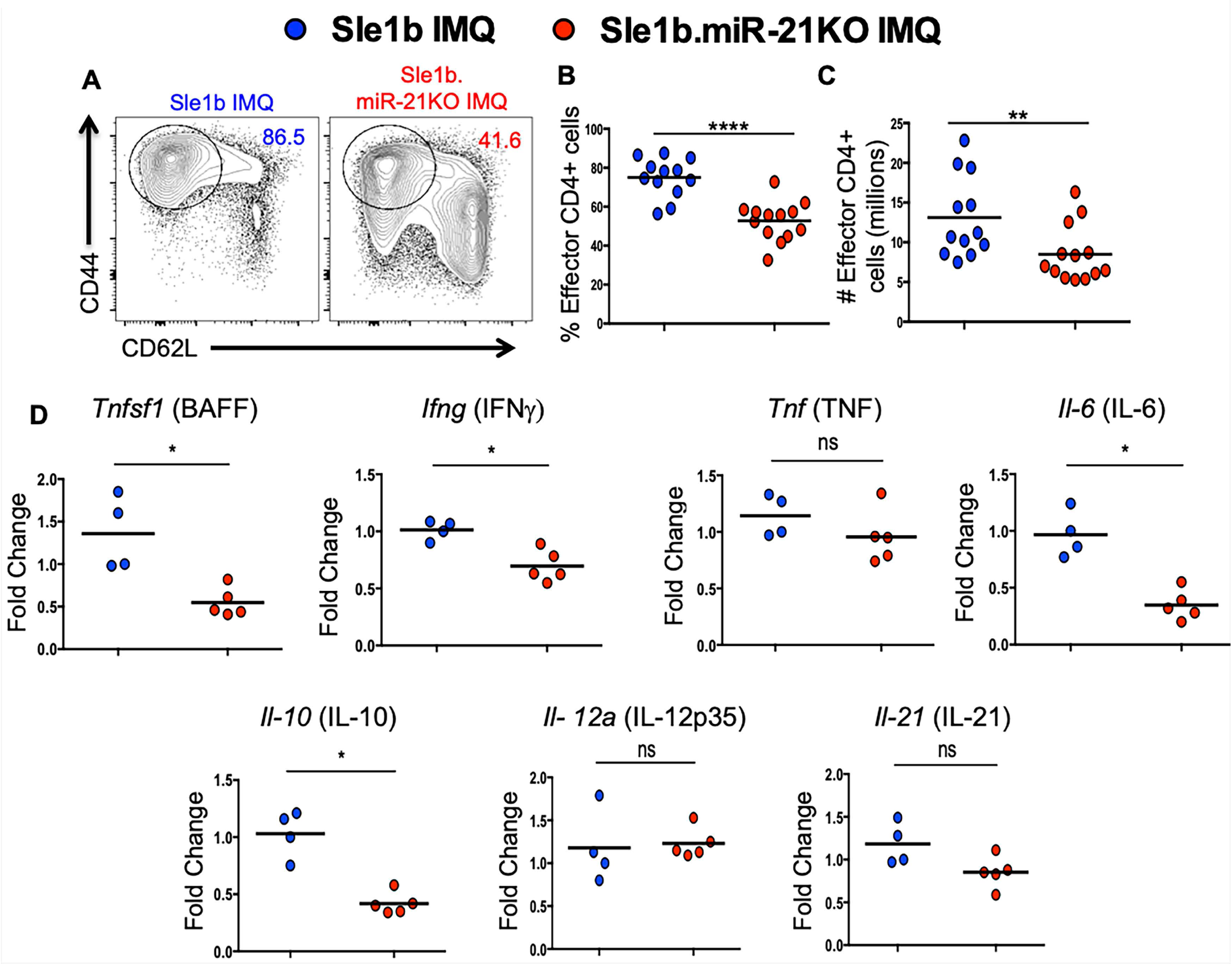
miR-21 Drives Effector CD4^+^ T Cell Responses and SLE-Associated Cytokine Production in the Spleens of Sle1b Mice. All data are from *Sle1b* and *Sle1b*.miR-21KO mice following 8-12 weeks of IMQ treatment from 2 independent experiments. Each data point represents one mouse. Blue circles are *Sle1b* IMQ and red circles are *Sle1b*.miR-21KO IMQ. **(A)** Representative flow plots of the effector CD4 T cell response (CD44^hi^CD62L^lo^). These plots are pre-gated on total CD4 T cells. **(B-C)** Quantification of the **(B)** percentage and **(C)** number of effector CD4 T cells within the total CD4 T cell population. **(D)** qPCR for the expression of BAFF, IFNγ, TNF, IL-6, IL-10, IL-12, and IL-21 in total splenocytes. Expression is normalized to HPRT. Two group comparison was performed by Mann-Whitney analysis. *p < 0.05, **p < 0.01, ***p < 0.001, and ****p < 0.0001.

### Cytokine Response and B Cell Activation Pathways are Altered in Sle1b B Cells in the Absence of miR-21

To determine which pathways and biological processes were significantly affected in *Sle1b* B cells in the absence of miR-21, we performed RNAseq analysis on B cells from mice of the *Sle1b* and *Sle1b*.miR-21KO genotypes following IMQ treatment. Analysis of B cells from IMQ-treated mice showed no significant differences in subpopulations of immature and mature B cells between the genotypes, although both strains had a similar reduction in MZ B cells (not shown). B cells derived from these strains of mice clustered separately by principal component analysis (PCA), indicating distinct transcriptional profiles in the presence or absence of miR-21 (Fig. 5A). Applying a p value cutoff of 0.05 and a fold change cutoff of 1.5, we found 195 significantly upregulated genes and 798 significantly downregulated genes in *Sle1b*.miR-21KO B cells. When applying stringent parameters with an adjusted p value (taking into account the chance of false discovery) of 0.05, we revealed a subset of 31 downregulated and top 10 upregulated genes in *Sle1b*.miR-21KO B cells compared to *Sle1b* B cells (Fig. 5B-C). The analysis of the top 31 downregulated genes predominantly revealed groups of genes that are involved in altered B cell response to interferon and cytokines, in addition to altered innate immune response (Fig. 5D), consistent with data that showed a difference in the expression of these factors in the spleens of these mice. We also performed gene set enrichment analysis (GSEA), which captures changes in the entire dataset. GSEA further indicated the enrichment of interferon and inflammatory cytokine response genes (Fig. 5E-G). GSEA also revealed an enrichment of genes involved in leukocyte differentiation and chemokine production (Figure 5K-L) and in cellular activation, proliferation, and apoptosis (Fig. 5H-J).

**Figure 5.**
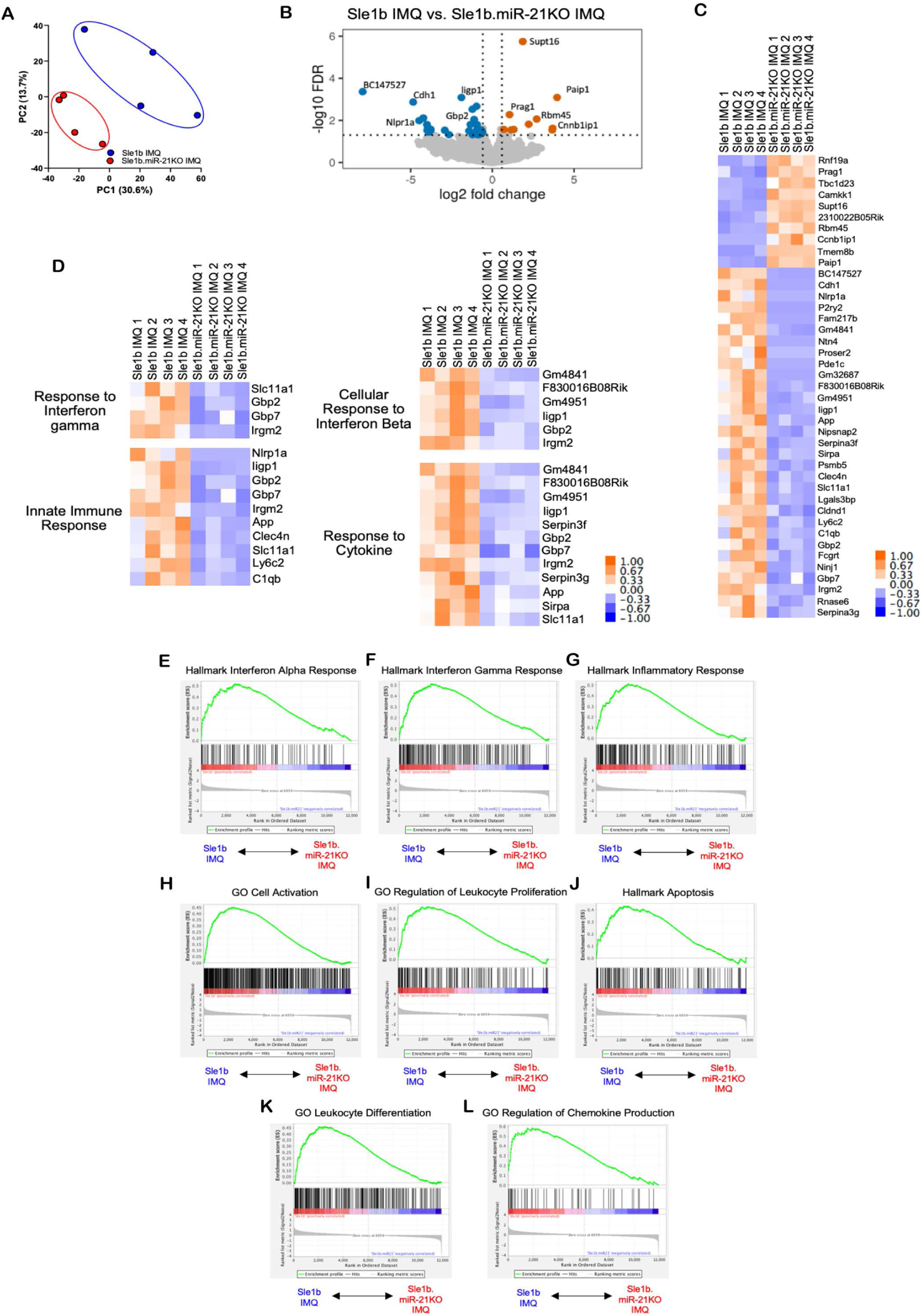
miR-21 Expression Alters Cytokine and Interferon Response and Activation Pathway in B Cells Derived from IMQ-Treated Sle1b Mice. Data are derived from sorted B cells from *Sle1b* and *Sle1b*.miR-21KO mice following 8-12 weeks of IMQ treatment. RNAseq was performed on B cells sorted from n=4 mice per group as indicated. Each sample represents B cells sorted from an individual mouse. **(A)** Principal component analysis (PCA) of the samples from each group (*Sle1b* IMQ-blue circles; *Sle1b*.miR-21KO IMQ-red circles). **(B)** Volcano plot indicating the differentially expressed genes from *Sle1b* and *Sle1b*.miR-21KO B cells that meet an adjusted p value cutoff of 0.05. Upregulated genes are depicted in red. Downregulated genes are depicted in blue. **(C)** Heatmap of the top differentially expressed genes between *Sle1b* and *Sle1b*.miR-21KO B cells that meet an adjusted p value cutoff of 0.05 and fold change of 1.5. Upregulated genes are depicted in red. Downregulated genes are depicted in blue. **(D)** Heatmap of the specific altered genes associated with select processes indicated by GO analysis. **(E-L)** Significant biological processes identified as enriched by GSEA (considering changes in the entire dataset). Additional information about generation of DEG lists and statistics can be found in the “Materials and Methods.”

GSEA analysis focusing on genes in intracellular signaling pathways identified the enrichment of genes in the p53, kRas, and PI3K/Akt activation pathways (Figure 6A-C). Consistent with differences in gene expression in these activation pathways, we found that B cells derived from *Sle1b*.miR-21KO mice had reduced expression of the activation markers CD80, CD86, and CD40 in vivo compared to *Sle1b* mice following IMQ treatment (Fig. 6D-I). These analyses support that miR-21 can alter B cell activation through a variety of mechanisms and pathways and supports the role of miR-21 in driving cytokine and interferon response in the spleen to alter B cell function.

**Figure 6.**
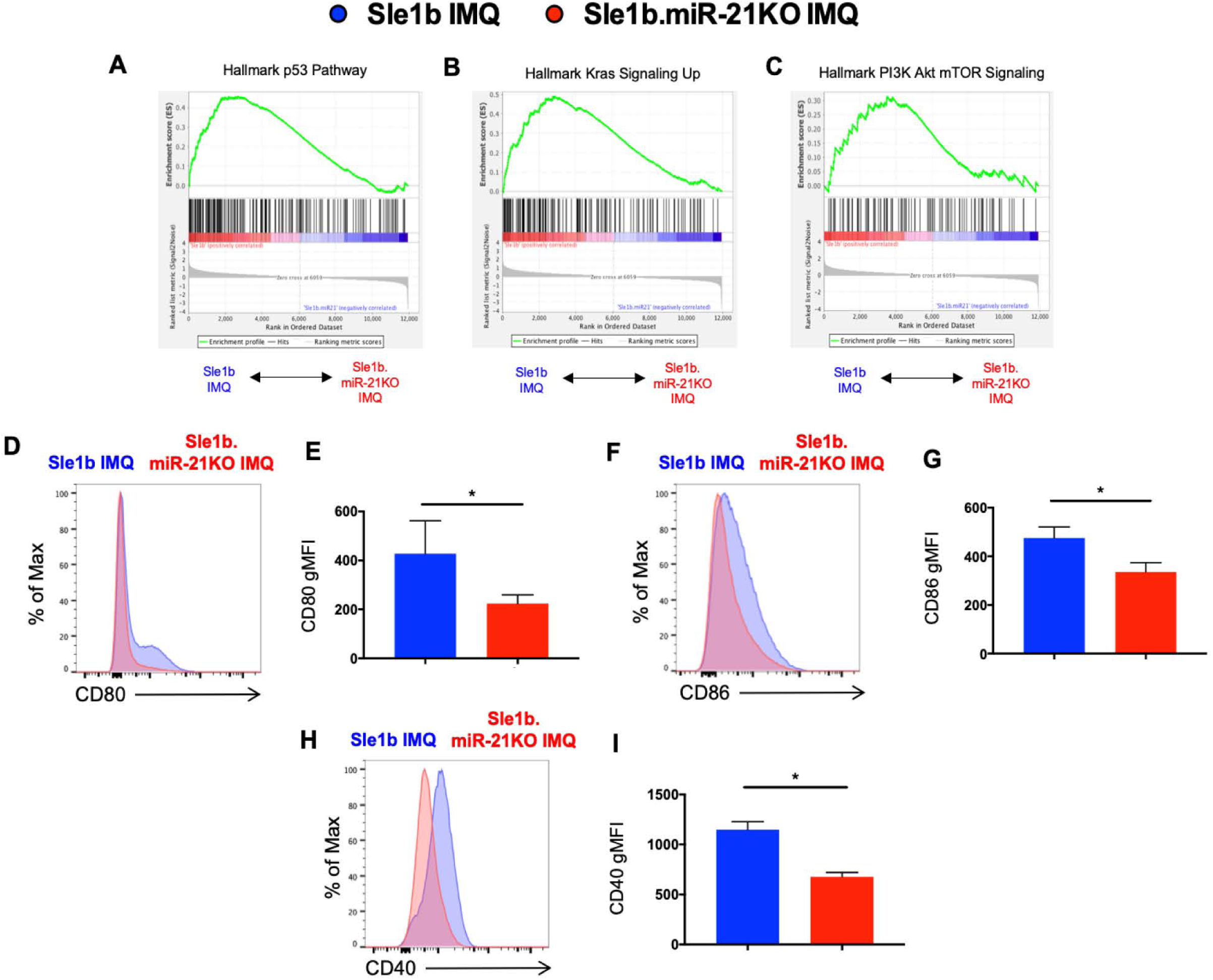
B Cell Activation Pathways and Activation are Driven by miR-21 in Sle1b Mice. Data are from *Sle1b* and *Sle1b*.miR-21KO mice following 8-12 weeks of IMQ treatment. Blue depicts *Sle1b* IMQ and red depicts *Sle1b*.miR-21KO IMQ. **(A-C)** GSEA plots for enriched genes in the **(A)** p53, **(B)** Kras and **(C)** PI3K/Akt/mTOR pathways. GSEA data is collected from the same mice and in the same manner as detailed in Figure 6. **(D, F, H)** Representative histograms showing the expression of **(D)** CD80, **(F)** CD86, and **(H)** CD40 on B cells derived from mice of the indicated genotypes. **(E, G, I)** Quantification of the expression of **(D)** CD80, **(G)** CD86, and **(I)** CD40 on B cells derived from n=4 and n=5 mice of the indicated genotypes. Bars on graphs represent the SD of the data. Two group comparison was performed by Mann-Whitney analysis. *p < 0.05, **p < 0.01, ***p < 0.001, and ****p < 0.0001.

### TLR7 and Type I Interferon Stimulation of Sle1b B Cells Results in miR-21 Upregulation and Modulation of miR-21 Target Gene Expression

We and others previously highlighted the critical B cell-intrinsic roles of TLR7 and TI-IFN signaling in SLE autoimmunity (6, 11, 14, 16). Further, our current RNAseq data indicated that cytokine and interferon-mediated activation of B cells from TLR7-stimulated *Sle1b* mice are dependent on miR-21. However, it remained unclear how miR-21 expression and miR-21 target gene expression in B cells may be regulated by TLR7 and TI-IFN stimulation. To study miR-21 expression and function, we profiled miR-21 expression in *Sle1b* B cells in response to TLR7 activation and TI-IFN stimulation and assessed the expression of confirmed and bioinformatically predicted target genes in stimulated *Sle1b* and *Sle1b*.miR-21KO B cells by qPCR array. miR-21 expression in *Sle1b* B cells was driven by R848, which was further amplified by the addition of IFNα4 (Fig. 7A). R848 and IFNβ dual stimulation drove the highest increase in miR-21 expression (Fig. 7A), which was maximal at 48hrs (Fig. 7B), thus miR-21 target gene expression was measured at this timepoint following stimulation. The miR-21 target gene array revealed that B cells express detectable levels of 68 select miR-21 target genes (Fig. 7C). Of these target genes, 26 were significantly upregulated in *Sle1b*.miR-21KO B cells (Fig. 7C-D; Table 1), indicating that miR-21 expression normally dampens their expression in *Sle1b* B cells in response to TLR7 activation and IFNβstimulation. Gene ontology (GO) analysis of these 26 genes revealed involvement in cellular activation, cell cycle, apoptosis and metabolic processes (Fig. 7E), which were also indicated during RNAseq analysis. These data collectively indicate that miR-21 expression in *Sle1b* B cells is enhanced by TLR7 activation and TI-IFN stimulation, which impacts the expression of miR-21 target genes involved in critical B cell processes.

**Figure 7.**
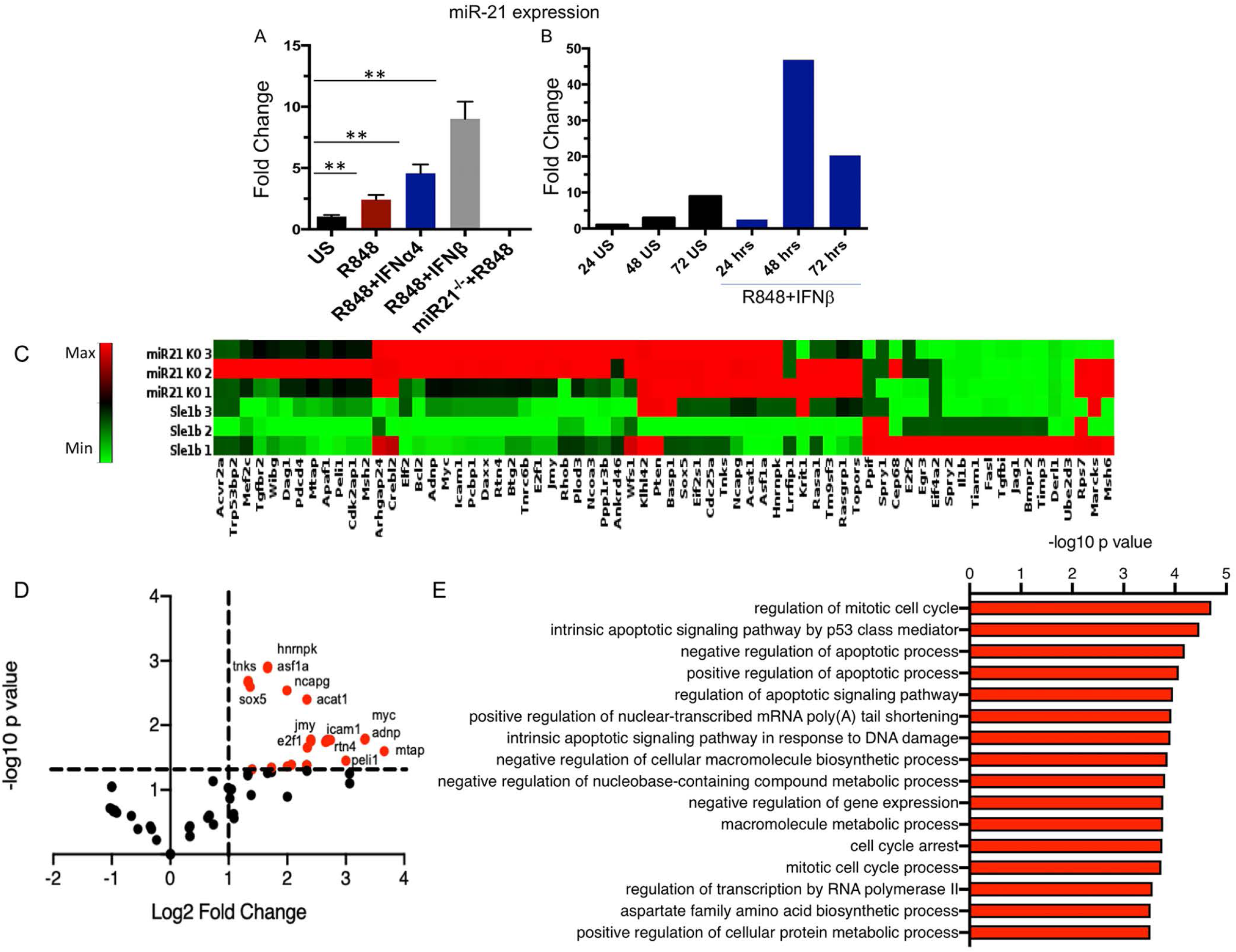
miR-21 Expression Increases in B Cells After IMQ and TI-IFN Stimulation, Leading to Suppression of Many miR-21 Target Genes. **(A)** Expression of miR-21 in *Sle1b* B cells following stimulation with R848, R848+IFNα4, or R848+IFNβ for 36 hours. **(B)** Timecourse of miR-21 expression in *Sle1b* B cells following stimulation with R848 and IFNβ. **(C)** Heatmap of miR-21 target gene expression in *Sle1b* and *Sle1b*.miR-21KO B cells via qPCR array following 48 hours of stimulation with IMQ and IFNβ. **(D)** Volcano plot showing the significantly upregulated genes in *Sle1b*.miR-21KO B cells from qPCR array analysis. Select genes are identified. **(E)** GO analysis of the 26 upregulated genes identified by qPCR array.

**Table 1.**
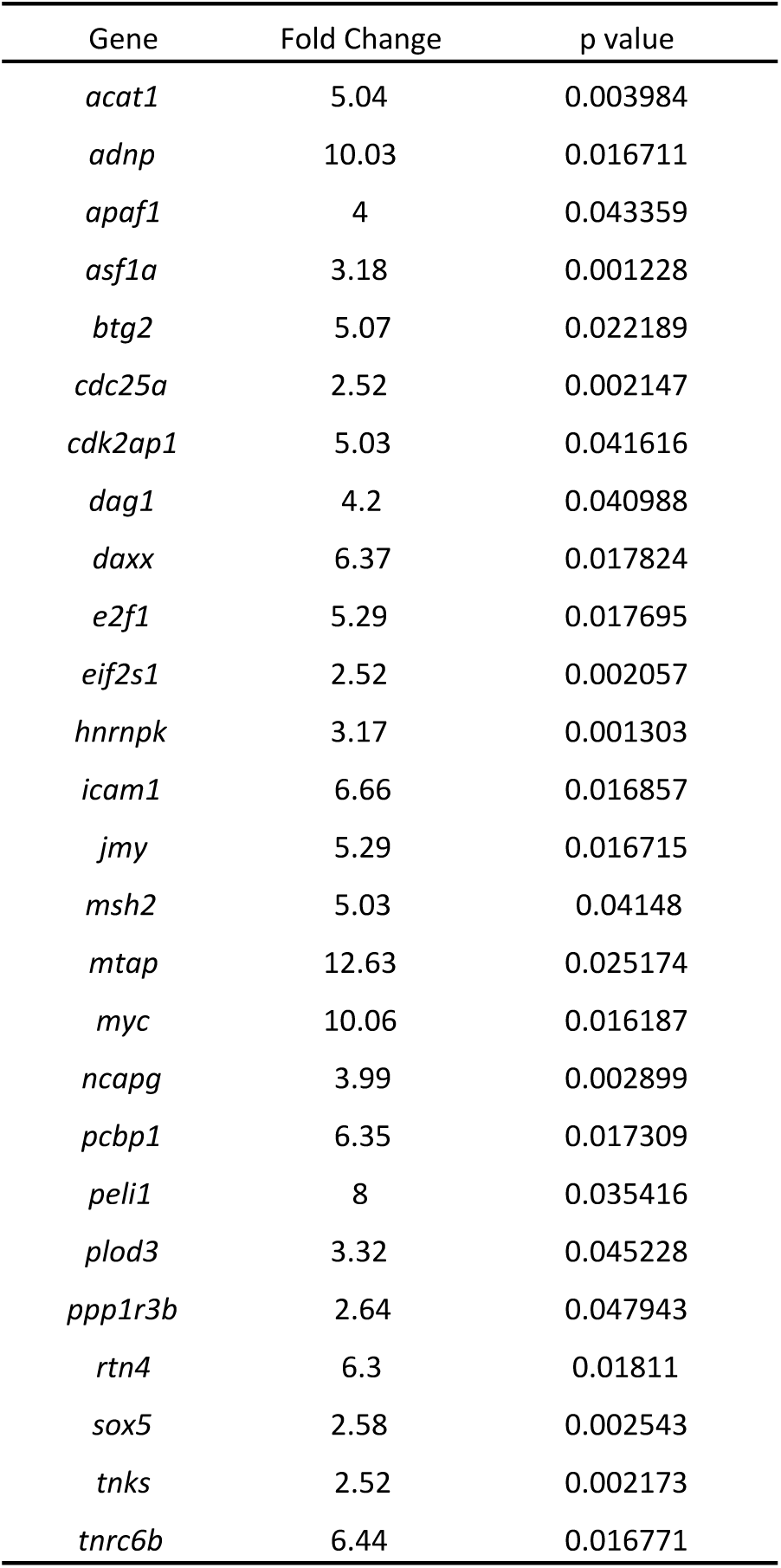
Upregulated miR-21 Target Genes In Stimulated *Sle1b*.miR-21KO B Cells

### miR-21 Regulates both Plasma Cell Formation and GC Responses in a TLR7-Overexpressing Spontaneous SLE Model

While we demonstrated a role for miR-21 in autoimmune AFC responses but not GC responses in the TLR7-induced IMQ SLE model, we wondered whether miR-21 exhibits similar activity in a separate TLR7-centric spontaneous autoimmune model, the B6.*Sle1b*.Yaa model. While both models focus on TLR7 contribution to autoimmunity, the models implement slightly divergent strategies. In the Yaa model, TLR7 is overexpressed whereas the IMQ system relies on the introduction of TLR7 ligand into the system to drive enhanced autoimmune manifestations. As such, we generated TLR7-overexpressing B6.*Sle1b*.Yaa mice deficient in miR-21 (designated *Sle1b*^Yaa^.miR-21KO). Analogous to IMQ treated *Sle1b*.miR-21KO mice, *Sle1b*^Yaa^.miR-21KO mice had reduced splenomegaly, and a reduced number of splenocytes and CD4^+^ effector T cells (Fig. 8A-E). Interestingly, in contrast to the IMQ model, the frequency and number of GC B cells and Tfh cells were significantly lower in *Sle1b*^Yaa^.miR-21KO mice compared to *Sle1b*^Yaa^ control mice (Fig. 8D, F). Consistent with flow cytometry data, histology showed a reduced frequency and size of GCs in *Sle1b*^Yaa^.miR-21KO mice (Fig. 8G, H). *Sle1b*^Yaa^.miR-21KO mice also had significantly reduced IgD^-^CD138^+^TACI^+^ plasma cell (PC) responses in the spleen (Fig. 8I) and a reduced number of autoantibody-producing AFCs in the spleen (Fig. 8J) and in the BM (Fig. 8K). These reduced responses in *Sle1b*^Yaa^.miR-21KO mice resulted in lower titers of serum IgG and IgG2c autoantibodies and reduced ANA seropositivity (Fig. 8L-N). We also analyzed innate cells in these mice and consistent with reduced splenomegaly we found numbers of innate cells including monocytes, moDCs and cDCs were much lower in the absence of miR-21 (Fig. 8O). Together, data from the spontaneous model indicate a role of miR-21 in regulating both AFC and GC responses, promoting autoantibody production and the development of SLE. These results further indicate that miR-21 control of autoimmune GC responses is sensitive to differences in autoimmune models, which may be linked to differences in the magnitude of activation that affect miR-21 activity.

**Figure 8.**
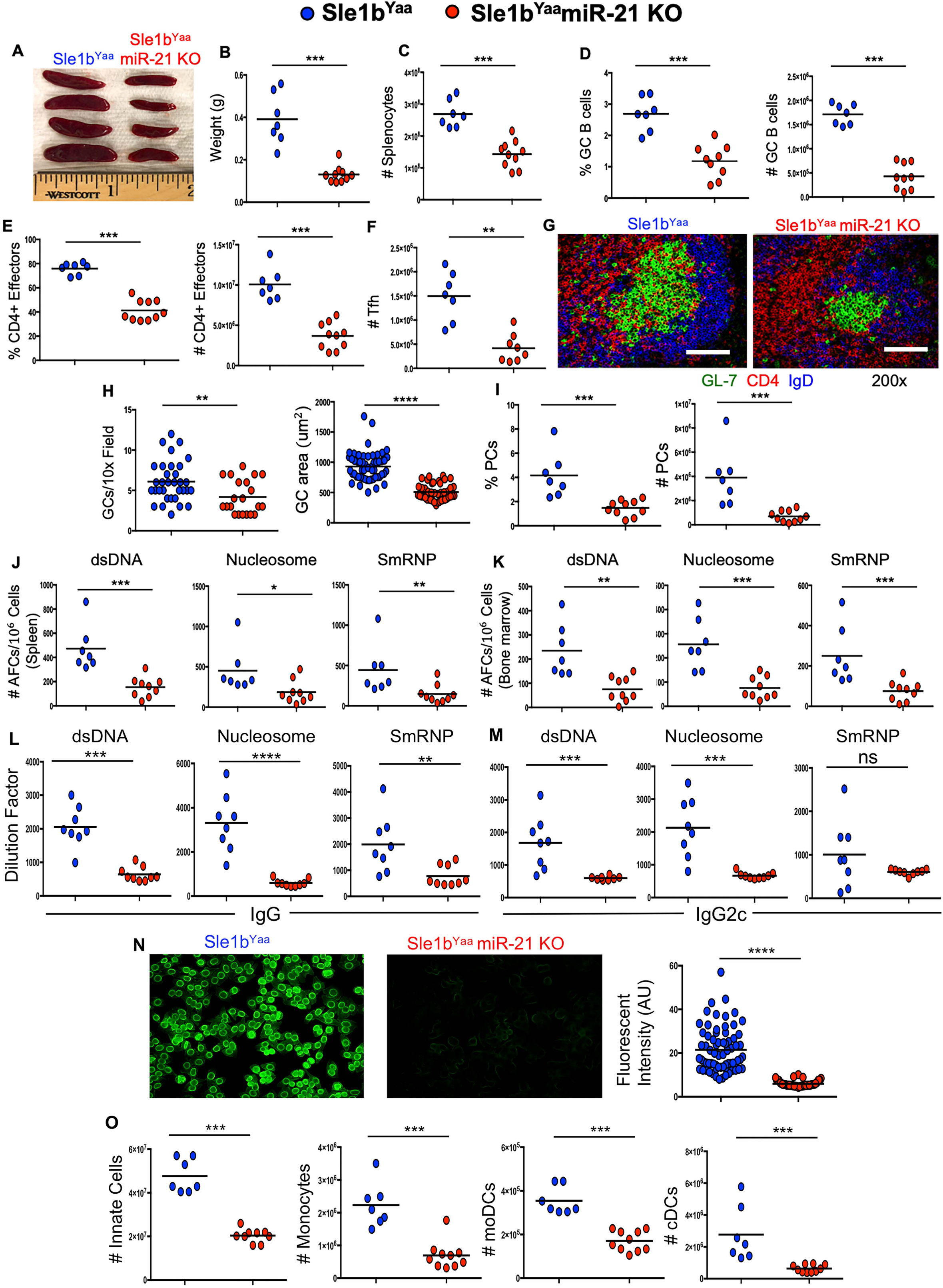
miR-21 Drives AFC and GC responses in TLR7-overexpressing SLE-prone mice. All data are from 3 mo old B6.*Sle1b*.Yaa (Sle1b^Yaa^) and *Sle1b^Yaa^*miR-21KO male mice. These data are from the analysis of spleens in 2 independent experiments, each with 4 or more mice per group. Each data point represents one mouse, except for HEp-2 quantitation which is detailed below. **(A-C)** Spleen size and weight, and quantification of splenocytes. (**D**) Flow cytometry analysis of the percentage and number of B220^+^GL-7^hi^Fas^hi^ GC B cells. (**E**) Flow cytometry quantification of the percentage and number of CD4^+^ effector T cells and (**F**) number of Tfh cells. (**G**) Histological analysis of the GC structure and (**H**) frequency and size. Scale bars in G represent 1001m. (**I**) Flow cytometry quantification of IgD^-^CD138^+^TACI^+^ plasma cells (PCs). Quantification of ELISpot analysis of autoantibody secreting B cells producing antibodies with reactivity against dsDNA, nucleosome and SmRNP in the (**J**) spleens and (**K**) bone marrow of the indicated genotypes of mice. Quantification and representative images of ELISpot analysis of autoantibody secreting B cells producing antibodies with reactivity against dsDNA and SmRNP in the bone marrow of the indicated genotypes of mice. (**L,M**) IgG and IgG2c serum titers of antibodies to dsDNA, nucleosome and SmRNP, and (**N**) ANA seropositivity in these mice. (**O**) Quantification of the numbers of innate cells and various myeloid cells such as monocytes, moDCs and cDCs. Two group comparison was performed by Mann-Whitney analysis. *p < 0.05, **p < 0.01, ***p < 0.001, and ****p < 0.0001.

### miR-21 is Required for Foreign Antigen Driven GC but not AFC Responses

As the contribution of miR-21 to autoimmune AFC and GC responses appeared to be model and context dependent, we further asked whether miR-21 was also required for B cell responses to foreign antigen, a separate system with different antigenic triggers. At 10d post-immunization with NP-KLH in CFA, we observed that the total number of B cells was reduced in the spleen of miR-21KO mice (Fig. 9A). We observed a significantly reduced GC B cell response in miR-21KO mice compared to WT mice, by total number and the percentage of B cells differentiating in this pathway (Fig. 9B-D). This was mirrored by a reduction in the follicular T cell response (Fig. 9F-H), although total number of CD4^+^ T cells were increased in miR-21KO mice (Fig. 9E). By staining with FoxP3, we further demonstrated that reduced percentage and number of follicular T cells in miR-21KO mice were caused by reduction in the number of both Tfh and Tfr cells (Fig. 9I-K). Consistent with the flow cytometry data, histology revealed smaller GC structures in miR-21KO mice than WT mice (Fig. 9L). Since immunization induces antigen-specific, class-switched IgG antibody responses, we assessed NP-specific antibody responses in the serum at 10d and 21d post-immunization. We found that NP-specific IgG and IgG1 responses were reduced two-fold in miR-21KO mice at 10d post-immunization, whereas NP-specific IgG2a/c was virtually absent in immunized miR-21KO mice (Fig. 9M). At 21d post-immunization, we found that NP-specific IgG1 and IgG2a/c responses were continually reduced in the serum (Fig. 9N). These results indicated an additional effect of miR-21 on GC and antibody responses in a separate system. We, however, did not observe significant differences in AFC responses, total CD4^+^ effector T cell responses, or in the innate cell responses (other than moDCs) during immunization (data not shown). Overall, miR-21 exhibits context dependent effects across multiple autoimmune and foreign antigen triggered responses.

**Figure 9.**
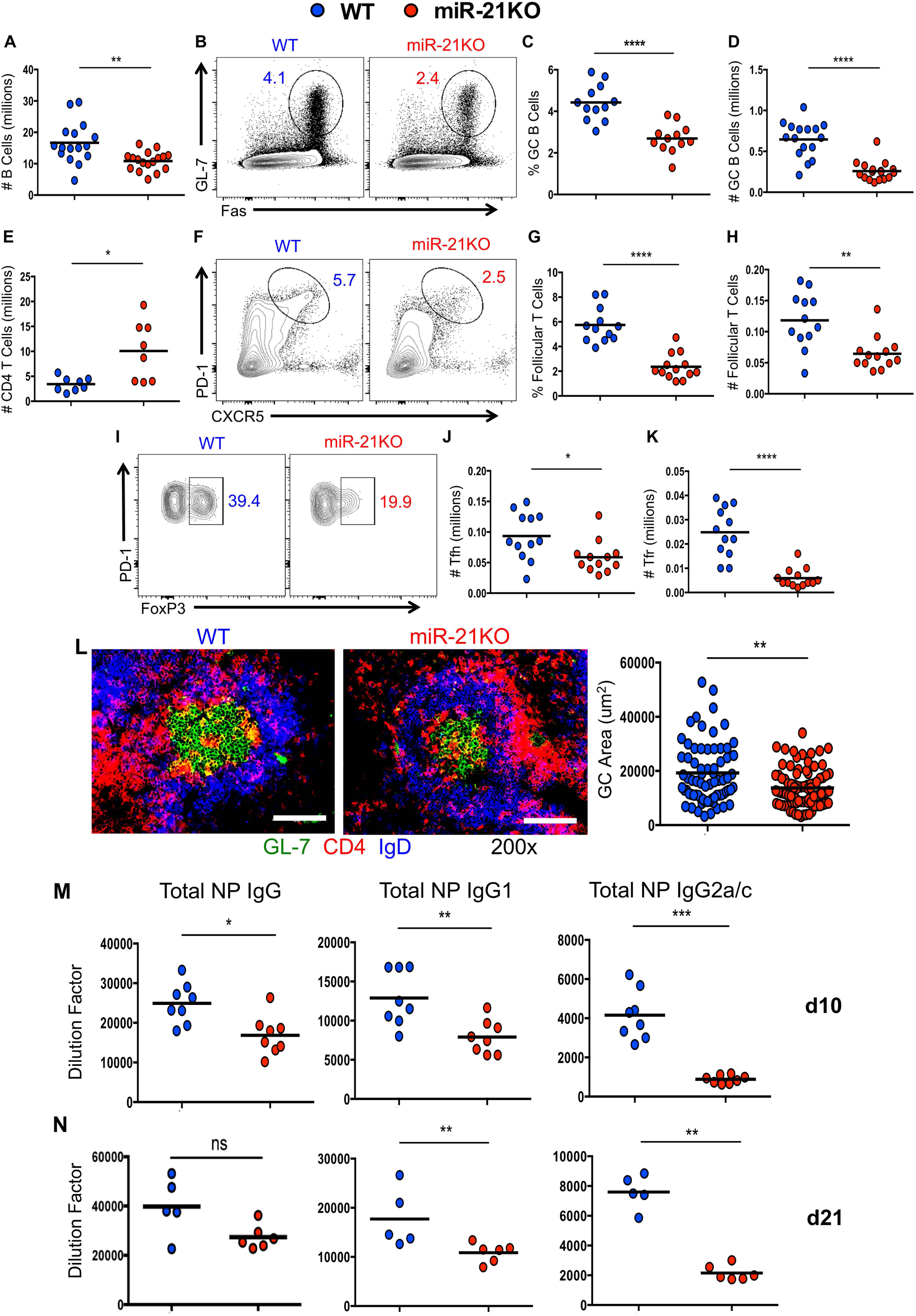
miR-21 Promotes TD-Ag Driven GC Responses. All data presented in panels A-L are from WT and miR-21KO mice at d10 post-immunization from 3 independent experiments with 3-5 mice per group. Except for histology, each data point represents one mouse (biological replicate). **(A)** Quantification of the number of B cells in the spleen. **(B)** Representative flow plots of the GC B cell (B220^+^GL-7^hi^Fas^hi^) response. **(C-D)** Quantification of the **(C)** percentage GC B cells within the total B220^+^ population and the **(D)** total number of GC B cells. **(E)** Quantification of the number of CD4 T cells in the spleen. **(F)** Representative flow plots of the follicular CD4 T cell (CXCR5^hi^PD-1^hi^) response. **(G-H)** Quantification of the **(G)** percentage follicular CD4 T cells within the total CD4 T cell population and the **(H)** total number of follicular CD4 T cells. (**I**) Representative flow plots for FoxP3^-^ Tfh and FoxP3^+^ Tfr cells within the total follicular CD4 T cell population. Total numbers of (**J**) Tfh and (**K**) Tfr cells. **(L)** Representative images of spleen sections stained for GL-7 (green), CD4 (red), and IgD (blue) and quantification of GC responses by measuring the GC area. Scale bars represent 1001m. Data represent at least 7 mice per genotype. **(M-N)** Serum antibody titers of NP-specific IgG, IgG1 and IgG2a/c at (**M**) 10d and (**N**) 21d post-immunization with NP-KLH. Two group comparison was performed by Mann-Whitney analysis. ns, not significant, *p < 0.05, **p < 0.01, ***p < 0.001, and ****p < 0.0001.

## DISCUSSION

The function of miR-21 in the loss of peripheral B cell tolerance and autoreactive B cell development, GC response, and AFC generation in SLE models remained unclear. Here using a TLR7-induced SLE model, we found that miR-21 has multifaceted effects on immune cells involving the modulation of B cell, T cell, and myeloid cell responses that may collectively promote a permissive environment for the generation of autoreactive B cells. Differences in the B cell response in this model were primarily observed in regard to the generation of plasma cell responses. Further, we observed that TLR7-mediated interferon and cytokine responses that drive SLE were reduced in the spleens of IMQ-treated *Sle1b*.miR-21KO mice. In addition, our RNAseq analysis of B cells from IMQ-treated *Sle1b*.miR-21KO mice showed a reduction in multiple interferon and cytokine activation signatures. Further, miR-21 target gene array analysis in B cells following stimulation with TLR7 agonist and IFNβ revealed increased miR-21 expression and modulation of many miR-21 target genes involved in activation, proliferation, and survival in *Sle1b* B cells. Notably, our data from the B6.*Sle1b*.Yaa spontaneous model highlighted the role of miR-21 in promoting autoimmune GC and Tfh responses in addition to recapitulation of its role in regulating several autoimmune features observed in the TLR7-induced model. Additional features observed in this model may be caused by differences in the overall magnitude of activation and miR-21 activity. Future study is warranted to distinguish if alterations in the cytokine and interferon signature are tied to reduced DC and monocyte infiltration or reduced T cell activation in these mice. Further, B cell intrinsic effects can be investigated using appropriate systems as miR-21 target arrays indicate that many previously identified miR-21 target genes (in other systems) may have a significant impact on B cell function in SLE.

Interestingly, we also found that the miR-21 is involved in TD-foreign antigen-driven B cell responses where it regulates the GC and antigen-specific Ab responses, but not the overall AFC response. Mechanisms underlying the dissociation of the Ab response with the AFC response currently remain unclear. These data indicate multifaceted and context-dependent roles of miR-21 in regulating autoimmune and foreign antigen driven AFC and GC responses, promoting both protective and pathogenic Ab production. This is likely dependent on the pathways and cell types activated in each independent model, as well as the magnitude of the response, which will impact the expression levels of miR-21 and its target genes during the various processes of B cell maturation and selection.

miRNA function as a whole in SLE has recently received attention, with important functions elucidated for multiple miRNAs (26-30, 49, 50). In regard to miR-21, increased expression in both mouse models and in humans had been associated with autoimmunity (32–37). miR-21 antagonism with a miR-21 sponge reduced splenomegaly and blood urea nitrogen levels in B6.Sle1.2.3 mice, though no extensive cellular characterization or mechanism was reported to shape future directed studies (32). Our current study using the TLR7-induced and spontaneous TLR7-overexpressing SLE models identified the cell types that are modulated during miR-21 mediated splenomegaly, loss of B cell tolerance and autoreactive B cell differentiation in two separate models. Under the heightened responses observed in TLR7-induced SLE model, miR-21 expression dysregulated splenic B cell selection events and plasma cell generation. Loss of tolerance through the GC versus extrafollicular pathway is unclear in this model, as selection events typically occur through both pathways. Regardless, reduced plasma cell generation in IMQ treated *Sle1b*.miR-21KO mice likely leads to the reduced long-lived autoreactive AFC responses observed. However, under chronic SLE-like autoimmune responses observed in the TLR7-promoted spontaneous B6.*Sle1b*.Yaa model, the GC pathway was significantly affected by miR-21 expression where miR-21 also plays an important role in autoreactive AFC generation, in addition to similar effects observed on plasma cell generation. Together, these data indicate a differential regulation of the GC response by miR-21 depending on the magnitude of the response. As such, this study adequately informs future comprehensive cell-intrinsic analysis that accounts for the potential diversity of miR-21 function among various immune cell types and in different SLE models.

Specifically, due to the alterations we observed in the B cell, T cell, and myeloid cell compartments in the spleen and the potential function of miR-21 in each of these cell-types, miR-21 could function through various mechanisms in multiple immune cell-types during the establishment and progression of SLE. Possible B cell-intrinsic effects of miR-21 are supported by the identification of 26 target genes in response to TLR7 activation and IFNβ stimulation by qPCR array analysis. Further investigation of the known functions of these 26 genes reveals sets of genes involved in the negative regulation of proliferation, activation, and survival, supporting the idea that miR-21 likely modulates these pathways to drive survival and proliferation of autoreactive B cells during SLE development and progression. As many genes are modulated, there is likely an additive effect of this modulation on function, rather than one specific target gene that is modulated to drive loss of tolerance alone. Ultimately, modulation of these genes may help autoreactive B cells to overcome an activation threshold that would otherwise not be met (51).

In addition to alterations in the B cell response, miR-21 deficiency in *Sle1b* mice also resulted in a reduction in innate cell numbers, including DCs, and effector CD4^+^ T cell responses in the spleen, which likely contributes to differences in B cell activation. Ultimately, these deficits may be caused by miR-21 effects on the myeloid cell compartment or in T cells, directly affecting their recruitment, activation, or polarization. Accordingly, previous study supports the requirement for miR-21 in the generation of myeloid precursors in the bone marrow during late stage sepsis (52). Further, miR-21KO mice have reduced monocytes in the blood (53), indicating a potential deficiency in production. These studies collectively indicate that miR-21 is involved in the recruitment of monocytes and DCs into the secondary lymphoid organs during TLR7-promoted autoimmune responses. Delineation of the source of cytokine and interferon, which can alter autoreactive B cell selection and survival (12, 13, 15, 16, 54), is of interest. Previous studies have indicated that intrinsic miR-21 expression alters the cytokine profile of macrophages, DCs, and T cells, including production of IL-6, IL-10, and IFNγ (55–60), which are reduced in IMQ-treated *Sle1b*.miR-21KO mice. Our previous study has indicated that CD4^+^ T cells are the major source of IFNγ in IMQ-treated *Sle1b* mice (40), indicating that miR-21 may affect the generation of Th1 cells in this system. As such, miR-21 is known to modulate T cell responses intrinsically (61), but further analysis is required to determine if effector CD4^+^ T cell responses are reduced in a miR-21 intrinsic or extrinsic manner. The specific subsets of immune cells that produce IL-6, BAFF, and IL-10 in a miR-21 dependent manner in our model of SLE requires further investigation, but may be tied to either reduced monocyte and DC responses or reduced effector CD4^+^ T cell activation.

Regardless of the intrinsic or extrinsic effects of miR-21 activity, our data indicate that miR-21 expression may promote the activity of several important characterized pathways in B cells, which may ultimately lead to a break in B cell tolerance and autoreactive B cell development in the AFC or GC pathway. IFN and cytokine receptor signaling can synergize with BCR and PRR signaling to activate signaling pathways such as PI3K and MAPK (62) pathways that were indicated by GSEA. MAPK signaling drives B cell proliferation and can promote autoimmunity (63–65). PI3K signaling is involved in B cell selection events and dysregulation of PI3K signaling in B cells also causes autoimmunity (66, 67). The interferons and cytokines that activate these pathways have been strongly tied to the loss of tolerance in SLE, further indicating that activation of these pathways is critical during loss of tolerance (12, 13, 15, 48).

The investigation into the role of miR-21 in the foreign antigen-induced GC response adds to the growing field of miRNA characterization in this microenvironment. miR-155 and miR-146 have been studied extensively and have roles in promoting and restraining GC responses in both T and B cell intrinsic manners (19, 21, 22, 68). Additionally, Tfh differentiation is dependent on the miR-17∼92 cluster (69, 70). Adding to these findings, we discovered a role for miR-21 in promoting the total magnitude of the GC, Tfh and antibody responses and in IgG2a/c class-switching following challenge with foreign antigen. As opposed to TLR7-driven autoimmune responses, we did not observe any effect of miR-21 on splenomegaly, extrafollicular AFC/plasma cell responses and an expansion of innate cell number upon immunization, albeit miR-21KO mice had reduced number of moDCs. It is currently unclear if there is a difference in the number of plasma cells reaching the bone marrow even at an early time point post-immunization, or if other mechanisms may be involved in the reduced antibody responses observed at d10 post-immunization. Assessing the autoimmune and foreign antigen responses together, these data indicate miR-21 mediated differential regulation of autoimmune and foreign antigen driven AFC and GC responses as well as expansion of innate and effector CD4^+^ T cells. While our immunization system has provided initial characterization, analysis of miR-21 in the context of pathogenic infection will provide insight into the requirement for miR-21 in generating protective antibody responses. This is of interest since viral infections polarize toward IgG2a/c production (71), which is compromised in miR-21KO mice following immunization.

In conclusion, our results highlight the effects of miR-21 in the loss of B cell tolerance and autoreactive B cell development, GC responses, and AFC development depending on the magnitude of SLE-like autoimmune responses, as well as the effects of miR-21 on the GC response during foreign antigen immunization, which represents a distinct mode of B cell activation and response. miR-21 may employ multiple mechanisms of action in various immune cell-types that exhibit altered response in its absence. Further analysis of the timing of miR-21 activity in SLE development and progression will also provide useful context into the temporal activity of miR-21 in these processes. Altogether, our data indicate a potential multifaceted and context-dependent miR-21 regulation of autoimmune and foreign antigen driven AFC and GC responses, which encourages further careful delineation of the underlying cell-intrinsic mechanisms.

## Abbreviations

Ab: antibody
AFC: antibody forming cell
ANA: anti-nuclear antibody
BCR: B cell receptor
GC: germinal center
IMQ: imiquimod
GO: gene ontology
GSEA: gene set enrichment analysis
miR: microRNA
SLE: systemic lupus erythematosus
SmRNP: Smith antigen/ribonucleoprotein
Tfh: T follicular helper cell
Tfr: T follicular regulatory cell
TI-IFN: Type I interferon

## ACKNOWLEDGMENTS

We thank Dr. Chetna Soni for insightful discussions. We thank the Pennsylvania State University Hershey Medical Center (PSUHMC) flow cytometry core facility for their assistance. We thank the PSUHMC Department of Comparative Medicine for animal housing and care. We also thank the Genomics Core Laboratory of Benaroya Research Institute for performing RNA sequencing and assisting in data analysis.

This work was supported by National Institutes of Health R21AI128111 to Z.S.M.R. and lupus foundation preceptorship to K.N.B, and the Finkelstein Memorial awards to both S.L.S and K.N.B.

